# Discovery and analysis of an 841 kbp phage genome: the largest known to date

**DOI:** 10.1101/2025.01.14.633092

**Authors:** LinXing Chen

**Affiliations:** The Department of Environmental Science and Engineering, University of Science and Technology of China, Hefei, China, 230026

**Author notes:** Corresponding author LinXing Chen.

**Keywords:** Huge phages, viruses, protein structure, metagenomics, genome curation

## Abstract

Viruses represent the most abundant biological entities on Earth, with bacteriophages (or phages) specifically infecting bacteria. Typically, double-strand DNA phages possess genomes around 50 kilobase pairs (kbp) in size, although the largest known genome surpasses 735 kbp. This raises intriguing questions about the potential maximum size of phage genomes. In this study, we present the first phage genome over 800 kbp, named BF3, which was reconstructed from oil reservoir production water. BF3 encodes 1,164 protein-coding genes (code 11) and 46 tRNA genes, and is classified within the Caudoviricetes, though its bacterial host remains unidentified. We utilized ColabFold to predict the structures of 744 protein-coding genes with high confidence, finding that 395 and 591 of these genes corresponded with known structures in the Protein Data Bank and the Big Fantastic Virus Database (BFVD), respectively. Notably, 153 of BF3’s predicted structures exhibited no similarity to those catalogued in the BFVD, which is currently the most extensive viral protein structure database. This study not only expands our understanding of phage genome capacities but also underscores the need for specialized analytical tools and pipelines to investigate exceptionally large phages.

## Introduction

Viruses are the most abundant biological entities on Earth, ubiquitously distributed across diverse ecosystems and capable of infecting hosts from all three domains of life ^1–4^. Among these, bacteriophages (or phages) are viruses that specifically infect bacterial cells. Phages play vital roles in microbial communities by lysing their bacterial hosts, influencing bacterial metabolism via auxiliary metabolic genes (AMGs), and driving evolutionary processes via horizontal gene transfer ^1^.

Phages typically possess genomes around 50 kbp in length. However, those with genomes exceeding 200 kbp are classified as jumbo or huge phages ^5,6^. Smaller phages generally encode 40-50 genes associated with minimal representation of AMGs. In contrast, huge phages harbor genomes that encode hundreds of genes, including those previously unidentified for basic viral life strategies in smaller phages. This extensive genetic repertoire goes beyond the typical functions such as attachment, DNA injection, DNA replication, viral particle assembly, and host cell lysis, encompassing complex and previously uncharacterized metabolic pathways ^6–9^. The advanced tools and analysis pipelines of genome-resolved metagenomics have greatly facilitated the identification of huge phages in the past several years, prompting a reevaluation of the theoretical limits of phage genome complexity and size.

In this study, we report the first phage with a genome size of over 800 kbp (841,704 bp), detailing its basic genomic features, and protein contents via structure predictions and comparison. We also discuss potential future research directions in light of methodological advancements.

## Results and discussion

### Huge phages are widespread across diverse ecosystems, yet their cultivation remains elusive

In 2019, Devoto et al. reported the first phage genomes exceeding 500 kbp, with an average size of approximately 540 kbp. These phages, named LAK phages, infect Prevotella spp. in the gastrointestinal tracts of various animals, including humans, pigs, and baboons ^10^. In 2021, a notable LAK phage genome, measuring over 660 kbp, was reconstructed from a horse gut metagenome ^11^. The LAK phages, likely the sole known phage group with genomes surpassing 500 kbp that have identified bacterial hosts, were recently classified into a new order within Caudoviricetes, named “Grandevirales” ^12^. Despite these discoveries, no LAK phage isolate has yet been successfully cultured.

Since 2020, an increasing number of large phages with genomes larger than 600 kbp have been documented ^6,11,13–17^, the largest of which surpasses 735 kbp ^6^. These genomes were reconstructed using genome-resolved metagenomics from various habitats, including freshwater, soil, animal guts, marine environments, landfills, groundwater, and sediments (Figure 1). Notably, the largest isolated huge phage to date has a genome of only 497,513 bp ^18^, a record that has stood for nearly fifty years.

**Figure 1.**
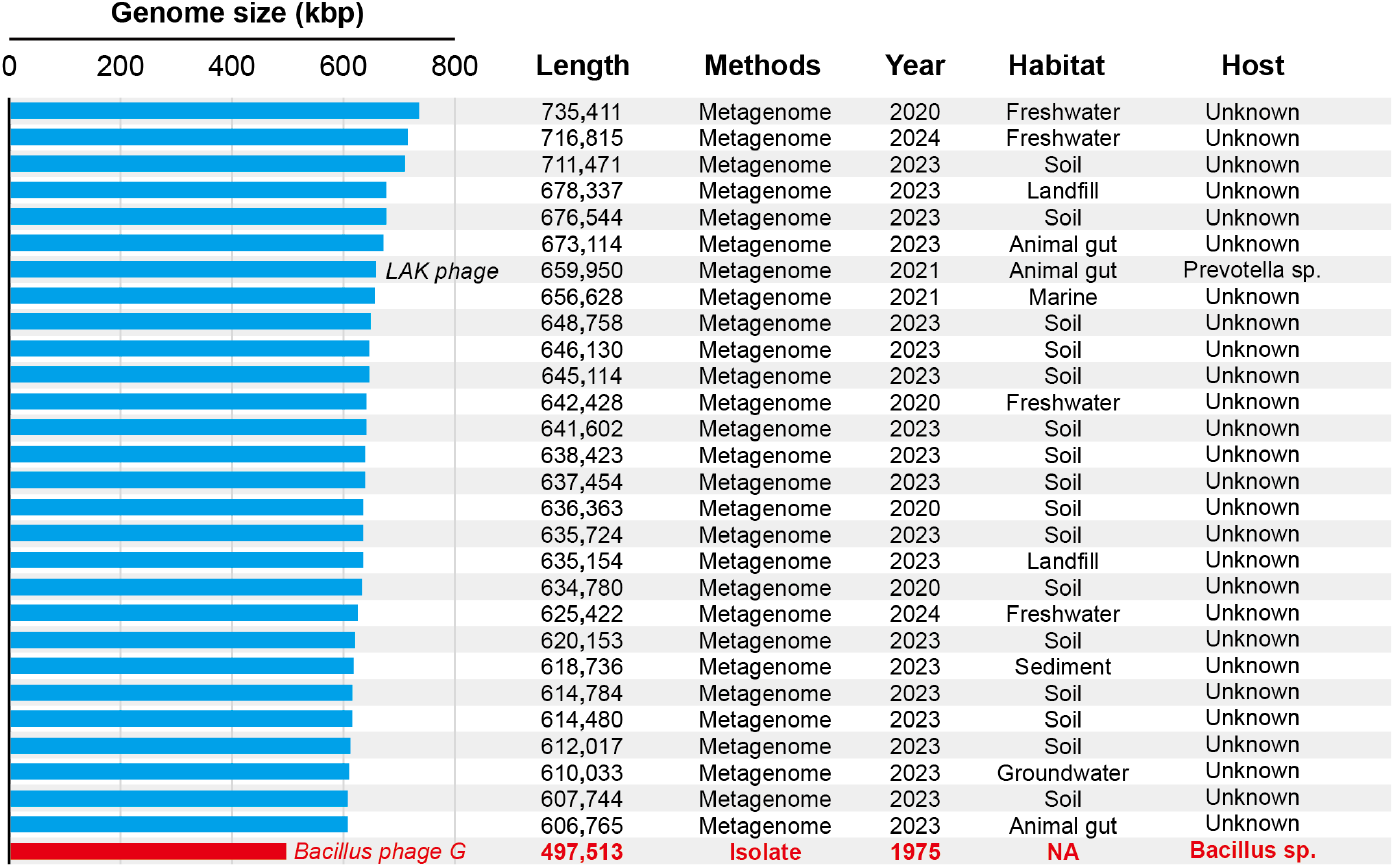
Summary of huge phages with genome size over 600 kbp. The isolated phage with the largest genome was included as a reference (highlighted in red). The information on methods, year, habitats, and potential bacterial hosts of the huge phages was retrieved from the corresponding publications and shown.

In summary, genome-resolved metagenomics has revealed that huge phages are broadly distributed across Earth’s ecosystems ^6^. Yet, the cultivation of these entities remains a rarity ^5^.

### The discovery and the feature of the largest phage genome known to date

An et al. ^19^ reported an incomplete phage genome with 708,116 base pairs (bp) in length. We attempted to curate the genome manually to see if we could obtain a phage genome exceeding 735 kbp, the longest reported so far ^6^ (see **Methods**). After several runs of reads mapping, genome extension, and assembly error fixation, we successfully obtained a circular phage genome with a length of 841,704 bp (Figure 2a). This genome was named BF3 hereafter, following the name of the sampling site ^19^. The mapping profiles of paired-end reads to BF3 with no mismatch allowed indicated no assembly error, no gap, and circular (Figure 2b, Supplementary Figure 1). However, various viral identification tools have different evaluation conclusions on this complete huge phage genome (Figure 2c).

**Figure 2.**
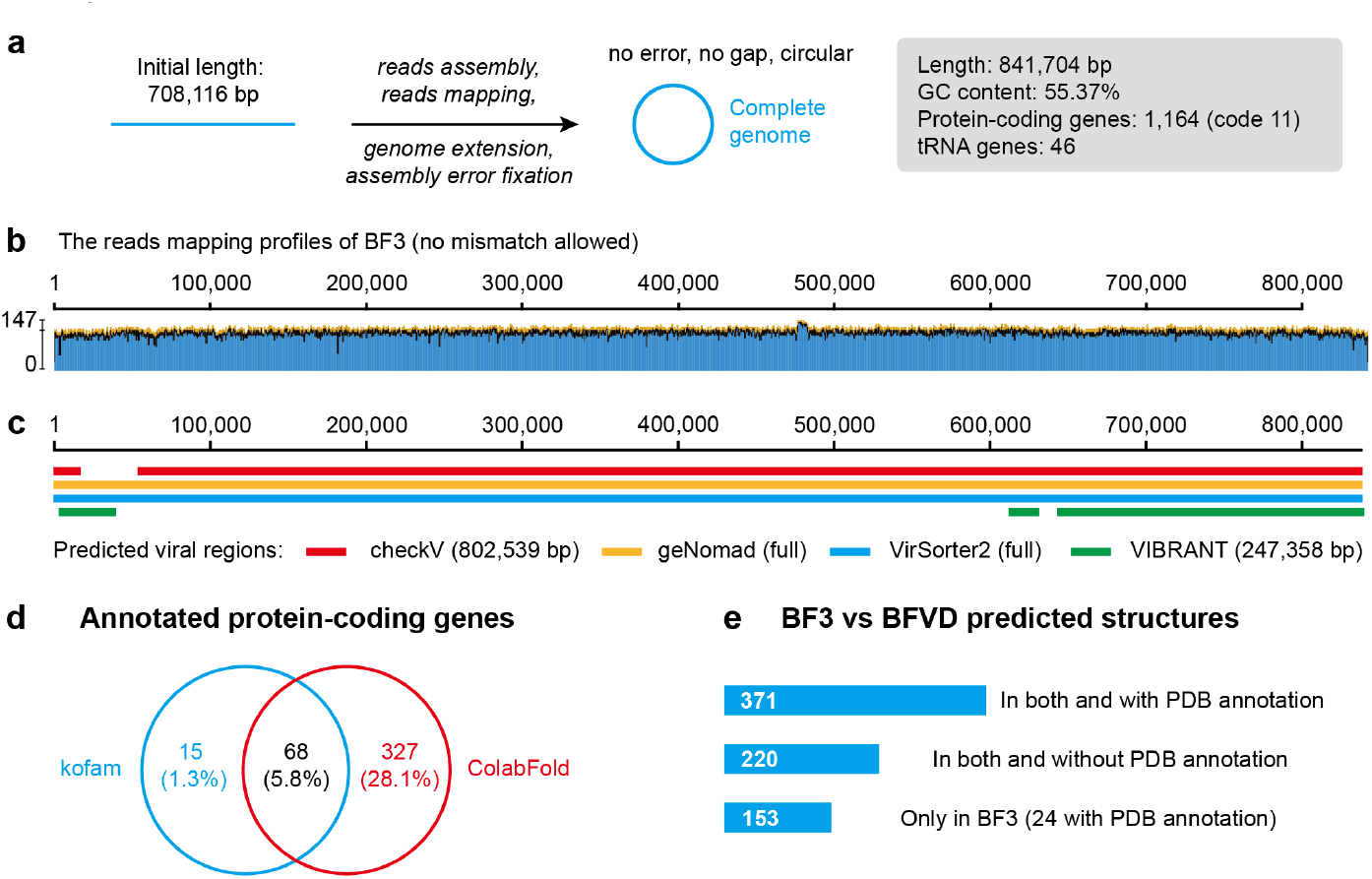
The metagenomic reconstruction and general features of the largest phage genome, BF3. (a) The brief analysis pipeline of reconstructing the complete genome of BF3, see details in the **Methods** section. (b) The reads mapping profiles of BF3. No mismatch was allowed for the mapping of paired-end reads. (c) The evaluation of BF3 by viral identification tools. (d) The number of protein-coding genes with known function annotated by kofam and ColabFold. The percentages were calculated according to the 1,164 protein-coding genes encoded by BF3. (e) The comparison of predicted protein structures between BF3 and BFVD.

BF3 is a member of Caudoviricetes as revealed by geNomad ^20^, and it has a GC content of 55.37% and encodes 46 tRNA genes. We predicted 1,164 protein-coding genes from BF3 using genetic code 11, with a coding density of 89.02%. Given that alternative coding may be applied, we tested all other available genetic codes and did not find any other one with a higher coding density.

We identified a total of 10,266 unique CRISPR-Cas spacer sequences from the assembled contigs. However, none of them had a significant match with BF3. Three *Maricaulis* spp.

(p Pseudomonadota;c_Alphaproteobacteria;o__aulobacterales;f_Maricaulaceae;g_Maricaulis) werepredicted by iPhop ^21^ to be the potential host of BF3. However, no *Maricaulis* spp. were identified in the corresponding community, though Alphaproteobacteria accounted for ∼50% of the community and members of Caulobacterales were with high relative abundance (Supplementary Figure 2). Thus, the bacterial host of BF3 remains unknown.

We annotated the protein-coding genes using kofam ^22^, and found only 7.1% of them (83 out of 1164) were with known functions (Figure 2d). To better understand the metabolic potential of BF3, we predicted the protein structures of the protein-coding genes using ColabFold (see **Methods**). As a result, 63.9% (744 out of 1164) of the proteins were annotated with reliable structure predictions (pLDDT ≥ 70), and 395 of the predicted structures have significant targets in the protein data bank.

We also compared the 744 reliable structures against BFVD ^23^, the most comprehensive viral protein structure database. We found that 591 of the structures had significant targets in BFVD, while BF3 also had 153 specific proteins, and 24 of them could be annotated by protein bank data (Figure 2e, Supplementary Table 1).

### Future research directions of huge phages

The discovery of phages with extremely large genomes opens new avenues for virology; however, innovative approaches are needed to explore these biological enigmas further. Firstly, to enhance our understanding and cataloging of huge phage genomes, developing new tools and analytical workflows are essential. These tools should be optimized to handle large datasets and complex genome assemblies typical of huge phages, improving accuracy and efficiency in identifying and assembling these large genomic sequences from environmental samples.

Secondly, cultivating and isolating phages with large genome sizes presents a significant challenge. Traditional methods have not yet yielded phage isolates with genomes surpassing 500 kbp ^18,24^. Therefore, there is a pressing need to innovate novel cultivation techniques or to modify existing methods to support the growth and study of larger phages. These advances would aid in understanding their life cycles and investigating their structural and functional characteristics.

Lastly, most genes encoded by huge phages are often not characterized through conventional sequence similarity searches, such as BLAST and HMM. The recent developments in structural prediction technologies, like those exemplified by AlphaFold ^25^ and its derivatives, such as ColabFold ^26^, offer promising pathways to build protein structure databases for these phages ^23,27,28^. Such a database would significantly enhance annotations and functional predictions of viral proteins. Furthermore, experimental approaches, including heterologous expression, gene silencing, or knockout studies ^29,30^, should be employed to validate the functions of genes of interest. This will allow a deeper exploration into the roles these genes play in the phage’s lifecycle and evolution, their interactions with host bacteria, and the broader environmental impacts.

These directions highlight the gaps in our understanding and chart a course for future research that could revolutionize our insights into phage biology, especially for those with unusually large genomes.

## Methods

### Huge phage genome collection

The published huge phage genomes > 600 kbp were collected from several publications ^6,11,13–17^. The downloaded genomes were clustered using the scripts (https://bitbucket.org/berkeleylab/checkv/src/master/scripts/) provided along with the CheckV tool ^31^ to remove redundant genomes (i.e., 100% similarity).

### Genome curation

The initial genome was published by An et al. ^19^, which was 708,116 base pairs (bp) in length and was not complete or circular (“BF3|k141_406419_flag=0_multi=53.6318_len=708116”, hereafter “BF3” for short). The paired-end reads of the corresponding metagenomic sample were downloaded from the China National Center for Bioinformation under the accession number CRR933431. The paired-end reads were *de novo* assembled using metaSPAdes v3.15.2 The paired-end reads were mapped to BF3 using Bowtie2 ^32^, and shrinksam (https://github.com/bcthomas/shrinksam) was used only to keep the mapped reads in the bam/sam file. The mapping profile was manually inspected using Geneious Prime version 2024.0.7 ^33^ to identify assembly errors, which were fixed as described previously^34^. The ends of the genome were extended using the consensus of unplaced reads mapped to the terminal regions, as detailed previously (https://ggkbase-help.berkeley.edu/genome_curation/scaffold-extension-and-gap-closing/). The extended regions of the genome were used to retrieve other contigs from the assembly. Once the retrieved contigs had similar sequencing coverage to BF3, they were assembled based on sequence overlap. Subsequently, the assembled genome will be mapped with the paired-end reads to confirm the assembly. Once confirmed, the consensus regions of unplaced reads mapped to the terminal regions will retrieve more contigs for assembly. Any assembly errors observed from the new contigs will be fixed as well. All these steps were performed again and again until there was evidence indicating that the genome was circular.

### Protein-coding gene and tRNA prediction

The protein-coding genes from BF3 and all assembled contigs ≥ 1000 bp were predicted using Prodigal V2.6.3 ^35^ (-m -p single) with the translation table 11. The tRNA genes were predicted using tRNAscan-SE 2.0.9 (July 2021) ^36^ with default parameters.

### Bacterial host prediction

Both the CRISPR-Cas spacer matching and iPhop ^21^ were applied to predict the bacterial host of BF3. For all the assembled contigs ≥ 1000 bp in length, the CRISPR repeat loci were predicted using pilercr v1.06 ^37^ using default parameters. Then, the protein-coding genes in the upstream and downstream 10,000 bp regions of the identified repeat loci were searched against the TIGRFAM HMM database ^38^ for Cas proteins. Only those spacers from the repeat loci with at least one Cas protein identified were extracted for target-matching analyses. The spacer matching analysis was performed using *blastn-short* with an e-value threshold of 1e-3. The iPhop analysis ^21^ was performed using default parameters.

### Community composition analyses

All the protein-coding genes predicted from contigs ≥ 1000 bp were searched against kofam ^22^ using kofam_scan-1.3.0. The hits of bacterial and archaeal ribosomal protein S3 (rpS3) (i.e., ’K02982’, ’K02984’) were retained and searched against the rpS3 proteins from GTDB genomes ^39^ using BLASTP ^40^. The taxonomy of the bacterial and archaeal rpS3 was assigned based on the top BLASTP hits. To calculate the relative abundance of each taxonomy, the sequencing depth of each rpS3-encoding contig was calculated using the jgi_summarize_bam_contig_depths script from metaBAT2 ^41^ from the mapping file, which was obtained by bowtie2 ^32^. For a given phylum, its relative abundance was calculated as the total sequencing depth of all rpS3-encoding contigs assigned to this phylum divided by the total sequencing depth of all rpS3-encoding contigs of the sample.

### Protein-coding gene annotation

The 1,164 predicted protein-coding genes were searched against the kofam database ^22^ using hmmsearch from HMMER 3.3.2 ^42^ with default parameters. The output was filtered to keep only the hits with an even or higher bit score, as provided by kofam.

### Protein structure prediction and comparison

The 1,164 predicted protein-coding genes were applied for ColabFold analyses, with the parameter set as “--num-recycle 3, --use-gpu-relax --amber”. The rank 001 models predicted structures with an average ColabFold pLDDT score of ≥ 70 (744 structures in total) were searched against the protein data bank ^43^ and the BFVD ^23^ using FoldSeek 9.427df8a ^44^ with the “easy-search” command (-c 0.5 --cov-mode 0). Only those with a TM score ≥ 0.5 (“alntmscore”) were thought to be reliable matches.

## Acknowledgements

This work was financially supported by the grant funding of KY2400000036. We thank the Super Computing Center at the University of Science and Technology of China for its support of ColabFold analyses.

## Author contributions

L.X.C. designed this study and performed all the analyses.

---

**Supplementary Figure 1.**
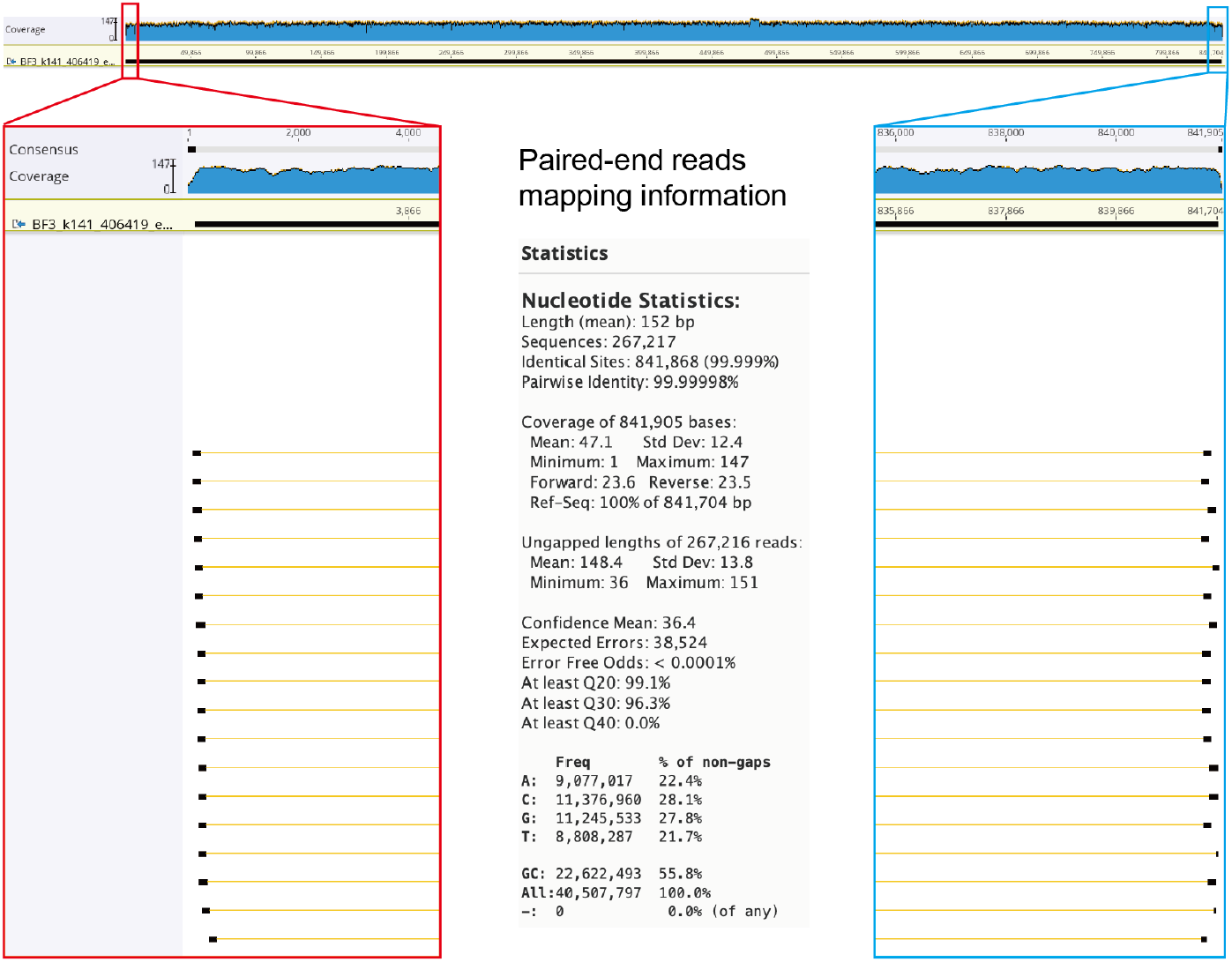
The paired-end reads mapping profiles of BF3. The zoom-ins show that paired-end reads span the two ends of the curated complete genomes. The mapping details are shown in the bottom middle.

**Supplementary Figure 2.**
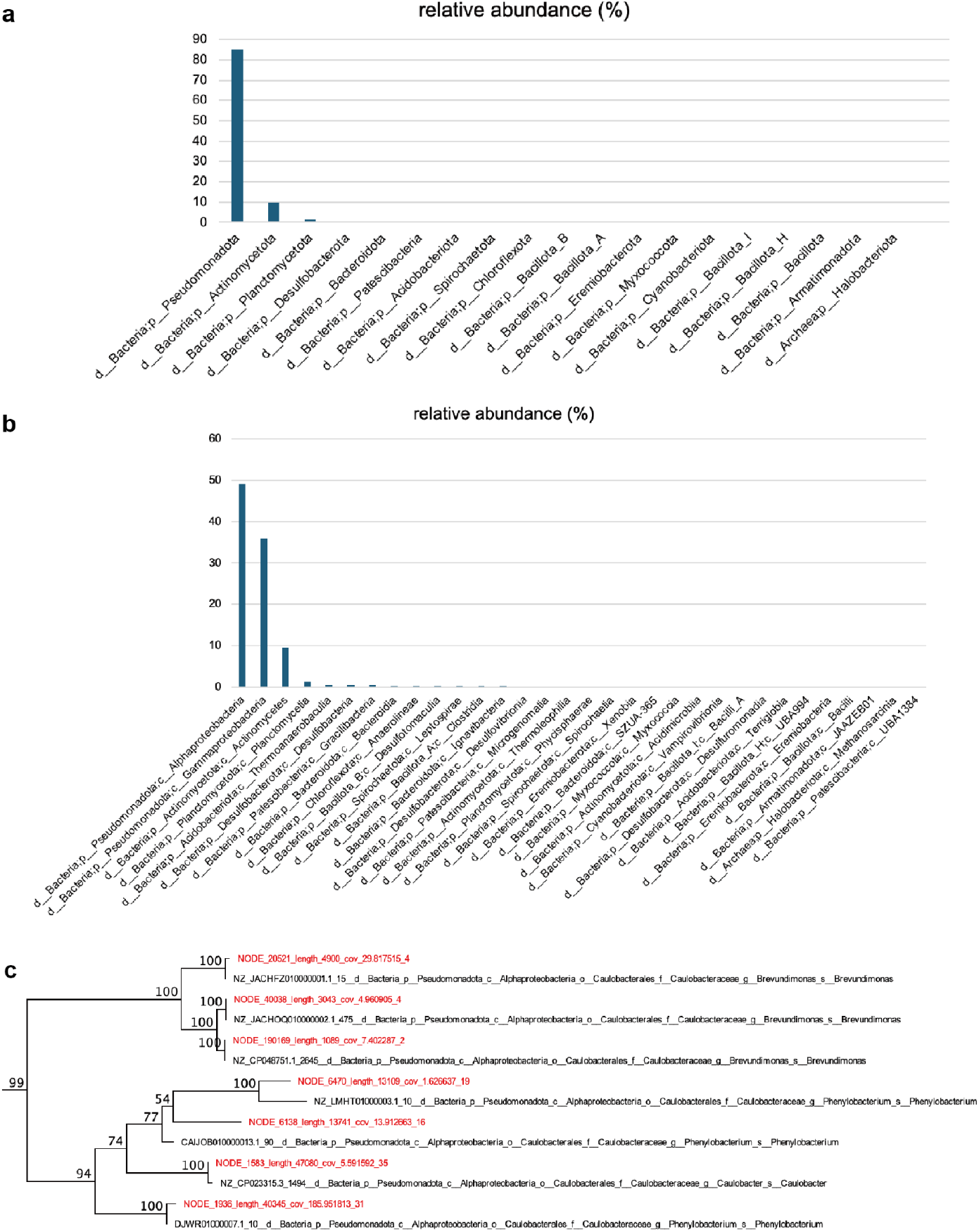
Alphaproteobacteria is the potential bacterial host of BF3. The community composition of the BF3 corresponding sample at the level of (a) phylum and (b) class. (c) The phylogeny of bacterial members (highlighted in red) from the order of Caulobacterales in the community. The numbers on the nodes indicated bootstraps.

